# FAST BOLD FMRI REVEALS THE SPATIOTEMPORAL COMPLEXITY OF NEUROVASCULAR COUPLING ALTERATIONS IN CEREBRAL SMALL VESSEL DISEASE

**DOI:** 10.64898/2026.06.14.732126

**Authors:** D Boido, Camélia Ressam, Valentine Perez, Benoît Beranger, Marianne Clary, Ali-Kémal Aydin, Jessica Lebenberg, Taleb Abbas, Fanny Fernandes, Denis Riviere, Zhong Yi Sun, Jean-François Mangin, Serge Charpak, Hugues Chabriat

## Abstract

Recent advances in neurophysiology highlighted the potential of high temporal resolution in Blood-Oxygen Level Dependent (BOLD) functional MRI (fMRI), although it is not yet standard practice. We demonstrated that fast BOLD fMRI can detect single-subject, single-stimulus visually evoked responses to brief stimuli at 3T. We used fast fMRI in patients with Cerebral Autosomal Dominant Arteriopathy with Subcortical Infarcts and Leukoencephalopathy (CADASIL), a genetic form of cerebral small vessel disease (cSVD). Using 2- and 10-second visual stimuli, we probed different neurovascular coupling regimes and showed that different regions of interest detect different facets of vascular dynamics. CADASIL patients showed significant changes in the amplitude and timing of the BOLD response, indicating early-age neurovascular impairment unrelated to anatomical lesions, and providing strong discriminative and generalization performance. These findings resolve prior inconsistencies in fMRI studies of CADASIL, supporting the use of fast fMRI to develop non-invasive biomarkers for cSVD and other neurodegenerative disorders.

## Introduction

Functional Magnetic Resonance Imaging (fMRI), particularly Blood Oxygen Level-Dependent (BOLD) fMRI^1^, ^1^has revolutionized the exploration of brain function. Since its first applications, this technique has demonstrated key advantages, including full-brain coverage and high spatial resolution, but also inherent limitations, most notably a restricted temporal resolution, which is fundamentally constrained by the hemodynamic nature of the signal. At the level of underlying mechanisms, neuronal electrical activity is mirrored by transient changes in the local blood flow, which ultimately corresponds to the broad definition of neurovascular coupling^2–4^. The local increase in blood flow, triggered by sensory stimulation or task-related increases in neuronal activity, constitutes the physiological response referred to as *functional hyperemia*,^5^ which is exploited by BOLD fMRI to map brain activation.

Biophysical modeling^6^ previously indicated that, at least, at low magnetic field (B_0_ <7T), the dynamics of BOLD functional hyperemia follows blood flow changes in veins, which result in an hemodynamic response function (HRF) peaking at 5 sec from the neuronal activation onset^7^. The intrinsic slowness of the HRF has traditionally constrained the temporal resolution of BOLD fMRI, typically imposing repetition times (TR) on the order of 1–2 seconds or longer. However, blood-flow-imaging techniques in mice using non-fMRI instruments, such as line-scanning two-photon microscopy or functional ultra-fast ultrasound, has revealed a faster coupling between neuronal activity and blood flow changes^8^, partially attributable to their higher temporal resolution (on the order of 200-300 ms). At high magnetic fields (7 Tesla and above), rodent BOLD fMRI studies have also shown HRFs faster than the 5-second peak reported in humans at low field strengths^9^. Such high-temporal resolution investigations have proven recently essential for addressing novel neurophysiological questions ^10,11^ (for a review, see^12^). Fast BOLD fMRI remains, however, rarely implemented in basic or translational research, reflecting the prevailing assumption that it offers limited added value relative to acquisitions performed at field strengths below 7 T.

Cerebral small vessel disease (cSVD) is a leading cause of stroke and dementia, yet the early detection of incipient vascular dysfunction remains highly challenging. Here, we used high-temporal-resolution BOLD fMRI at 3T to characterize cortical hemodynamic responses to 2- and 10-second visual stimuli in healthy controls and patients with CADASIL ^14,15^, the most common genetic form of cSVD. Our specific analysis revealed that CADASIL patients exhibited marked alterations in the amplitude and timing of the BOLD response, indicative of an early neurovascular impairment. These results not only help reconcile inconsistencies reported in previous fMRI studies of this condition^13,14^ but also pave the way for the development of non-invasive biomarkers of vascular dysfunction in cSVDs. They also provide evidence that fast BOLD fMRI, even at limited magnetic field strengths daily used in clinical practice, can reveal novel biomarkers of vascular dysfunction.

## Methods

### Ethical information

Written informed consent was obtained from all participants to the study. The study and consent documents were approved by the Comité de Protection des Personnes (CPP) Île-de-France IV (registered with the US Department of Health and Human Services, IRB 00003835) under the following protocol references: CPP 2018/98, Promoter 18.10.25.69641, and ID-RCB 2018-A02489-46.

### MRI acquisitions

All MRI data were acquired on a Siemens 3T Verio scanner with a 32-channel head coil. Anatomical 3D T1 whole brain scans were performed with a 1mm isotropic MP-RAGE. For functional acquisitions, our objective was to use a standard “clinical” 2D GRE-EPI sequence, yet with fine-tuned parameters to capture temporal details of the hemodynamic response. High temporal resolution of 300 ms was achieved using 12 slices of 3 mm thickness, with an in-plane resolution of 3 mm. A MultiBand= 3 acceleration factor was sufficient to achieve the 300 ms resolution target. The echo time was set to 30 ms, and the flip angle was 38°. The fMRI field of view was positioned to encompass the entire visual cortex, with the angle set using the medial border of the cerebellum as the reference point. Visual stimuli were programmed using Psychtoolbox, which runs in MATLAB. We used a 6 Hz flickering checkerboard. The contrast was set to 100%. A fixation cross was displayed when the stimulation was OFF. The 2-second stimuli were followed by 26 seconds of rest, while the 10-second stimuli had 50 seconds of rest to allow a complete recovery of the vascular response. In the middle of the 5’ sequence, i.e., after the 6^th^ 2-second stimulus and the 2^nd^ 10-second stimulus, the fixation cross turned red and was slightly enlarged to ask the subject to click a button to verify the wakefulness state. For one patient, the 10-second stimulation data could not be acquired due to a technical issue.

### MRI analysis

FMRI analyses have been conducted with SPM12 and other custom-made Matlab and Python scripts, and are presented in the schematics of Suppl. Fig. 3. Functional MRI data alignment on the T1-weighted anatomical acquisition has been verified and, if needed, manually performed with a rigid transformation. Movement correction was performed with SPM12 using minimal smoothing parameters. Noticeably, the high temporal resolution of our dataset allowed us to clearly rule out the displacement due to breathing and heartbeat, which were clearly visible in the movement regressors. No time correction was performed thanks to the high temporal resolution. We used Marsbar^15^ (SPM plugin) to extract the fMRI time course from the regions of interest (ROIs). The ROI of the Calcarine Sulcus was automatically segmented with BrainVISA Morphologist^16^ and manually separated from the Pericalcarine Sulcus. The correctness of such a segmentation was blindly verified by two experts in sulcal morphology (D.R. and J.-F. M). The Visual Cortex ROI used in our analysis was taken from the Cuneus region of the atlas by Shen and colleagues^17^. To preserve the fMRI from interpolation, we moved the atlas onto the anatomical acquisition of each subject. The t-test ROI was made by comparing 27 volumes (8.1 sec) before and during the activation brought by the 2-second or 10-second stimuli. This data-driven approach was preferred over a ‘standard’ General Linear Model to avoid making any assumptions about the shape of the hemodynamic response, which was potentially impacted by the diseases, as outlined in our working hypotheses. To merge the information from the four 5’-sequences recorded with 2-second stimulation, for each voxel, we averaged the p-values from each sequence.

In each individual, we also used a novel deep learning-based segmentation tool for measuring white matter hyperintensities on FLAIR images, the most consistent and prominent MRI feature of cerebral lesions accumulating since the early stage of the disease (htt ps://github.com/miac-research/MARS-WMH)^18^. The lesions masks were then corrected visually by a trained reader with a large experience in evaluating CADASIL MRI data. The volume of WMH was then normalized by the intracranial cavity volume measured using ANTS (https://github.com/ANTsX/ANTs).

### Features extraction

Feature extraction was conducted with custom Python scripts on BOLD responses, averaged across the voxels identified by the tested ROIs, and then averaged across stimulus repetitions. Before any signal analysis, the average BOLD fMRI response has been median-filtered with a kernel size equal to 7. The area under the curve (AUC) ratio was computed using the measure of the AUC in the beginning (between 3 and 6 sec, or 12 and 15 sec, for the 2 and 10-second stimulation respectively) and in the end of the fMRI responses (between 9 and 10.5 sec, or 16 and 18 sec, for the 2 and 10-second stimulation respectively). The response duration was measured by assessing the time point at which the return to baseline occurred. To avoid the confounding effect due to noisy fluctuations and prevent data loss due to responses never reaching the baseline during the allocated time window, we opted for a linear fit over the points between 6 and 17.5 seconds for the 2-second stimulations. For 10-second stimuli, instead, we had to accommodate the fitting time window for each subject to account for the higher variability of the decay phase of these responses. The response amplitude and time to peak were computed from the stimulus onset to the maximum of the average, median-filtered response. The time to half-peak was measured using the time value at which the amplitude was half of its maximal value. Similarly, the decay phase was assessed by measuring the time value at which the amplitude was half of its maximal value during the decay phase. We preferred this parameter to evaluate decay over other parameters, such as linear and logarithmic fits, because it was very efficient in highlighting the different decay dynamics, thereby avoiding any arbitrary choice of the fitting time window or function. The response shape was quantified using kurtosis and skewness, as well as the full-width at half-maximum (FWHM).

### Statistical analysis

We used *a priori* statistics, setting the confidence level at 95%; however, we reported higher confidence levels when reached, using the standard symbology with asterisks. Each statistical test used is reported in the manuscript, along with its p-value and degrees of freedom. Statistical tests between patients and controls were performed using a two-tailed parametric (Welch) t-test or a Mann-Whitney U-test when normality was not met (tested with the Shapiro normality test). Statistical tests have been performed in Python using the SciPy^19^ and Scikit-Learn^20^ libraries. No multiple comparison corrections have been applied.

### Data and code availability

The individual imaging data analysed in this study are not publicly available because they contain sensitive personal data and are protected under the European General Data Protection Regulation (GDPR) and relevant ethics approvals. Raw MRI data therefore cannot be shared openly. Non-identifiable, derived data supporting the findings of this study, including subject-level summary parameters extracted from averaged hemodynamic responses, are available from the corresponding author upon reasonable request and subject to appropriate data-sharing agreements. Note that the analysis pipeline is depicted in the Suppl. Fig. 3 and used standard SW for MRI analyses. Custom-made MATLAB and Python scripts have not been placed in a repository because they perform simple tasks described in the Methods and do not constitute substantial or original contributions to MRI data analysis. However, custom analysis scripts developed by the authors are available from the corresponding author upon request and can be shared with reviewers during peer review.

## Results

### 1. A fast BOLD fMRI protocol with two types of visual stimulation at 3T

We designed a visually stimulated fast fMRI protocol (TR = 300 ms) to trigger two vascular activation regimes: a brief vascular response to a 2-second stimulation and a sustained activation of the neurovascular unit to a 10-second stimulation (Fig. 1a). To set the time duration of these two stimuli, we analyzed a previous fMRI dataset^21^ with 20- and 40-second stimulation blocks and estimated the expected response amplitudes through convolution^22^. The predictions suggested that 2-second stimuli produce half of the maximal response, while 10-second stimuli are sufficient to obtain the maximal response (Suppl. Fig.1).

**Figure 1.**
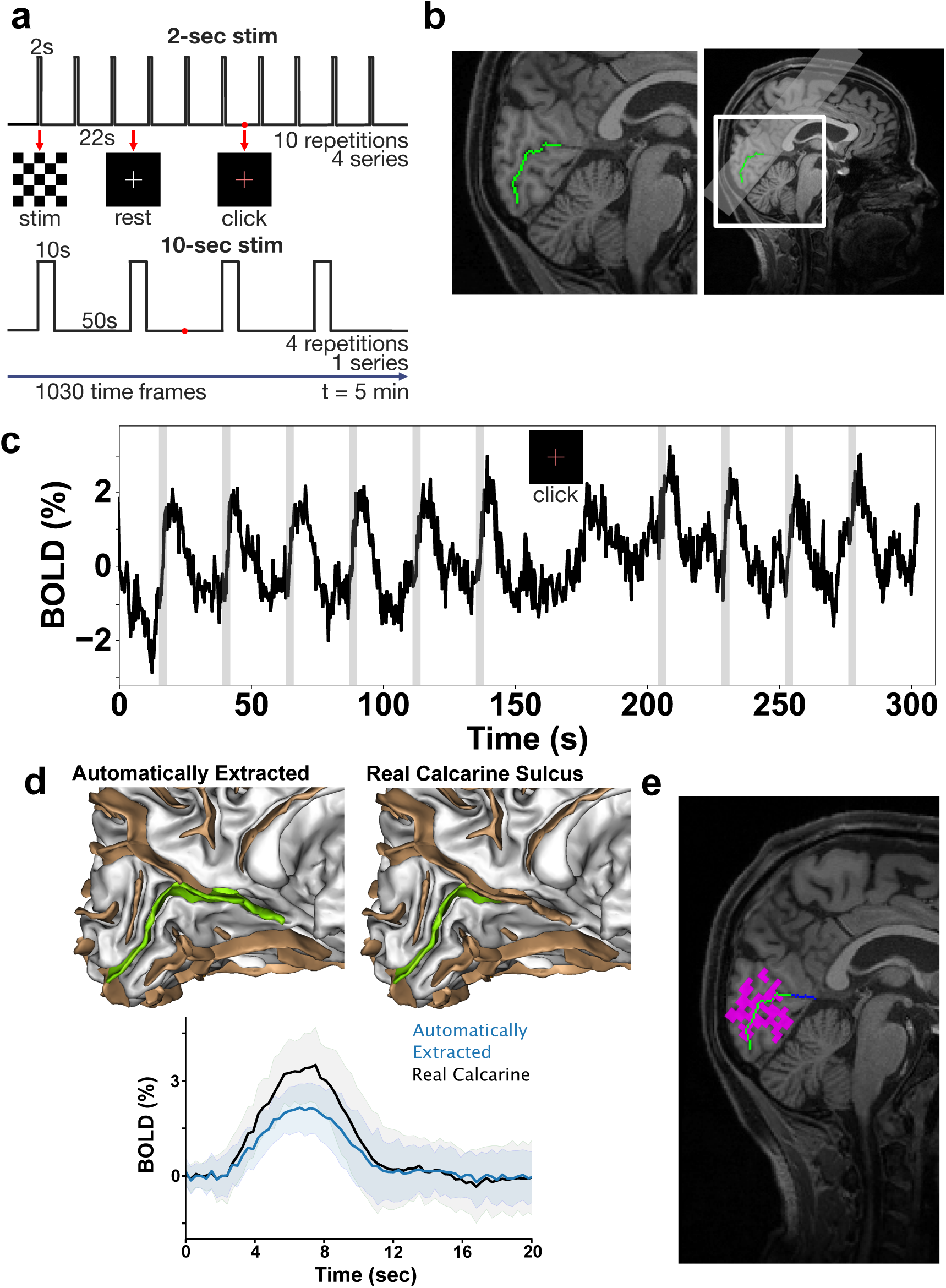
The fast fMRI protocol. (a) Schematic representation of the stimulation protocol, consisting of a sequence of 2 sec or 10 sec visual stimulations (flickering checkerboard). The short stimuli were repeated 10 times per sequence (4 series), whereas the long ones were repeated only 4 times, in a single series. Roughly in the middle of each sequence, the fixation cross turned red to require the subject to click a button to inform about wakefulness state. (b) A relatively small field of view covering the visual cortex was required to record at high temporal resolution. The Calcarine Sulcus is depicted in green. (c) BOLD signal time course of a series of 2 sec stimuli extracted from the Calcarine Sulcus. Each visually-evoked response is clearly visible. Notably, button-click also elicited a detectable BOLD signal in the visual cortex. (d) The AI-based algorithm provided by BrainVISa Morphologist cannot distinguish between the calcarine and the continuation of the occipital sulcus, so the calcarine was manually extracted for each processed subject (right panel). Segmentation refinement led to a more accurate evaluation of visually evoked BOLD responses (bottom panel). (e) As expected, the calcarine overlaps with the statistically activated voxels.

We thus implemented 4 sequences of 5 minutes for the 2 s stimulation to collect a total of 40 responses and a single sequence with 4 repetitions of 10 s stimuli. Subjects were asked to click a button during each 5-minute acquisition, upon displaying a red cross to ensure that they were actively engaged during the task (Fig. 1a). We decided to initially restrict our analysis to the calcarine sulcus, hereafter called “calcarine”, which is known to provide a high BOLD contrast and corresponds to a reliable region of interest (ROI) across subjects (Fig. 1b). With this fast fMRI sequence, after averaging functional voxels comprising the calcarine and a thin gray matter layer around it, we could clearly observe single visually evoked BOLD responses, as predicted by our simulation (Fig. 1c). Notably, clicking the button upon the display of the red cross triggered a large BOLD fluctuation detectable in the calcarine ROI, justifying the delay of 26 s that was inserted before the subsequent visual stimulus. We assessed test–retest reliability in three subjects scanned twice on the same day (Suppl. Fig. 2a) and observed very high reproducibility, with correlation coefficients of r = 0.87 ± 0.16 and r = 0.96 ± 0.03 for the 2- and 10-second stimuli, respectively (mean ± St.Dev, Pearson’s test). Using a sophisticated algorithm dedicated to the automatic extraction of cortical sulci (BRAINVISA Anatomist^16^), we were not able to separate the calcarine sulcus consistently from a nearby sulcus defined by the algorithm as peri-calcarine. As the functional activation of the peri-calcarine is not specifically related to the visual system, we performed an additional manual segmentation to isolate the calcarine sulci (Fig. 1d). Although this more stringent selection did not substantially alter the BOLD dynamics, a strong effect of this more stringent selection on the response amplitude was detected in 14 out of 15 subjects (Fig. 1d and Suppl. Fig. 2b). This refinement was further validated by t-test activation maps, which showed no meaningful activation in pericalcarine voxels (Fig. 1e).

Because fast BOLD fMRI sequences are rarely employed in clinics, and their integration with short visual stimuli is even less conventional, we aimed to determine how averaging across different ROIs affects the resulting BOLD visual response. Averaging over the real calcarine produced large single-stimulus BOLD responses in the range of 2-4%, as in the example time course shown in Fig.2a, illustrating the average responses of 40 repetitions on the right side. Averaging over the entire visual cortex still preserved a clear response to single visual stimuli (Fig. 2b, time course), but with a smaller amplitude (Fig. 2b, right). Using a data-driven voxel selection to define the ROI based on t-test activation maps—thus avoiding any a priori assumptions about the shape of the hemodynamic response function—yielded BOLD response amplitudes comparable to those obtained from the calcarine ROI, while substantially reducing noise (Fig. 2c). Despite differences in ROI definition, the average responses from the three ROIs showed comparable temporal dynamics.

**Figure 2.**
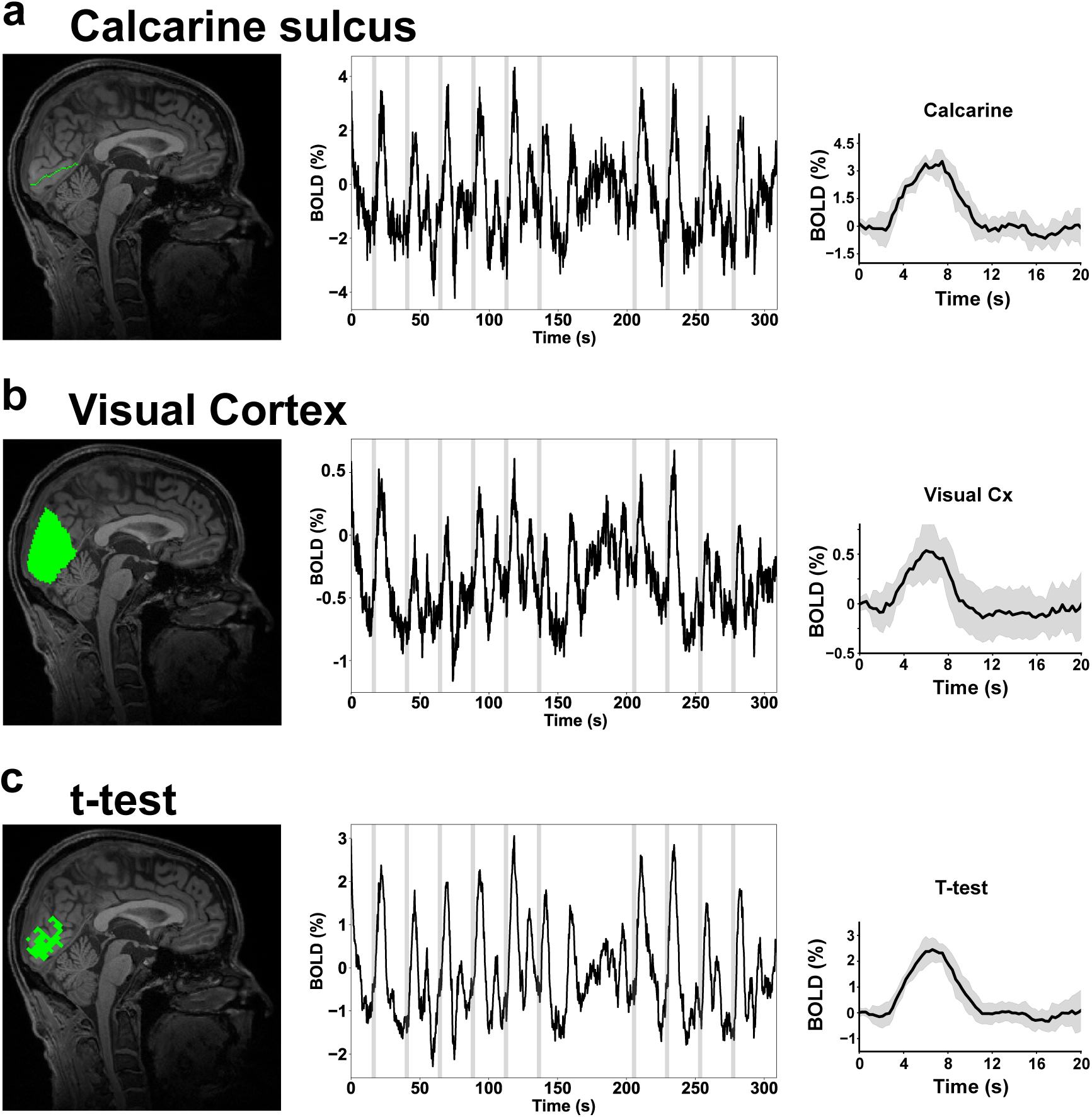
Impact of the region-of-interest on average fMRI response. Examples of BOLD time courses and average responses from different ROIs. (a) In the calcarine the signal is well detectable at single-stimulus level and average responses reach 2% amplitude. (b) Averaging across the entire visual cortex produces smaller responses usually around 1% amplitude. (c) Responses averaged across t-test selected voxels have comparable amplitude to calcarine but display lower fluctuations on time courses.

### 2. A targeted fast fMRI analysis for detecting specific disease alterations in CADASIL

We hypothesized that such a fast BOLD fMRI protocol could reveal subtle vascular dynamic alterations in pathological conditions where measurable changes are limited, and used CADASIL as a paradigmatic small vessel disease to explore this approach. This genetic disorder alters progressively the wall matrix of cerebral blood vessels, which presumably contributes, after decades, to the accumulation of ischaemic lesions and neurodegeneration. Prior fMRI investigations in patients with CADASIL have consistently shown no or limited BOLD dynamic alterations, in contrast to the results obtained in cerebral amyloid angiopathy^23^.

We then scanned a cohort of 18 patients with CADASIL (51 ± 11.31 years) and 16 age-matched control subjects (52 ± 9.14 years). It is important to note that we ensured that none of the patients aged between 32 and 71 years had any comorbidity that could affect the brain, and that none were taking medications known to alter vascular function. All of them were pauci-symptomatic, none exhibited any clinically significant or disabling cognitive or motor symptom. In the calcarine ROI, we observed that the BOLD fMRI responses to short visual stimuli were markedly slower in patients than in controls (Fig.3a). The difference became particularly apparent when the responses were normalized to their amplitude (Suppl. Fig. 4). To further characterize the apparently slower BOLD responses in patients, we quantified the kurtosis and skewness of their Gaussian-like distributions fitted to the response. Both parameters differed significantly between the two groups (Fig. 3b), indicating more asymmetrical and smoother response curves in patients with CADASIL than in controls. When the analysis was repeated using data-driven voxel selection from t-tests, the results were like those obtained in the calcarine. Analyses of time to peak and response duration yielded concordant results across both approaches, suggesting that the slowdown affects both the rising and decay phases of the response. (Fig. 2c). In contrast, the analysis of all visually evoked fMRI responses obtained in the entire visual cortex abolished all previous significant effects (skewness: p= 0.14; kurtosis: p= 0.06; duration: p= 0.15; time to peak: p= 0.11, t-tests).

**Figure 3.**
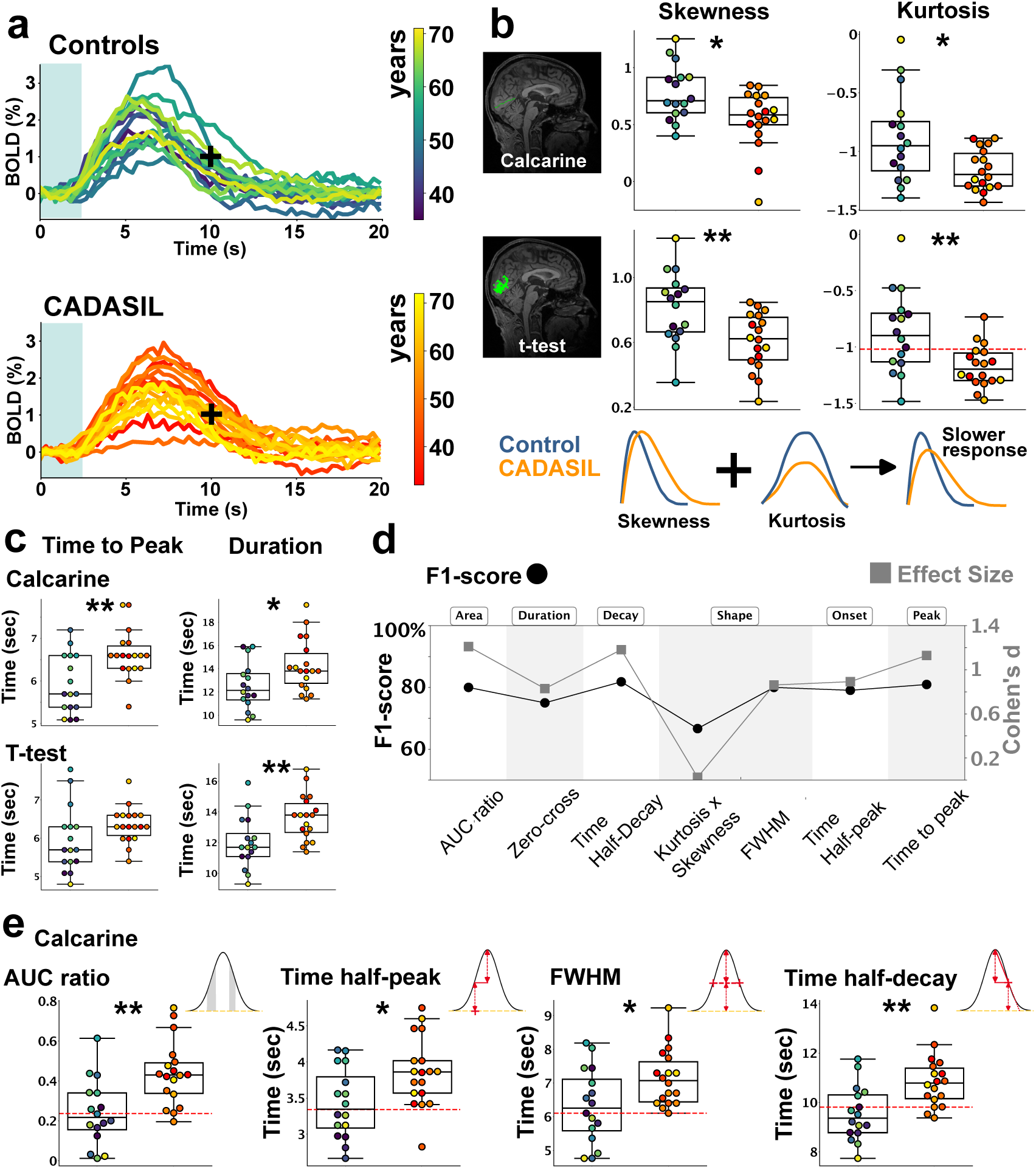
Short visual stimulation in CADASIL and Control subjects. (a) BOLD responses to 2 sec stimuli averaged in the calcarine for CADASIL and Control subjects, with color-coded subject’s age. (b) The quantification of responses’ skewness and kurtosis in calcarine shows a clear indication of slower responses in CADASIL than control subjects (skewness, p= 0.026, kurtosis; p= 0.018, t-test), confirmed by the average of the t-test selected voxels (skewness, p= 0.0055, kurtosis; p= 0.007, t-test). (c) Time to peak and duration of the BOLD response in calcarine and with t-test selection further confirmed the slowdown for CADASIL’s responses versus Controls (time to peak, p= 0.003, duration, p= 0.02, t-test). (d) F1-score and effect size values across 7 computed parameters accounting for different phases and features of the BOLD responses indicate a discrimination power in the order of 80%. Generalization assessment equal or over 1 (Cohen’s d) in 6 out of 7 parameters. (e) Selection of the best F1-score ranked parameters (AUC ratio, p= 0.001, time half peak, p= 0.01, FWHM, p= 0.02, time half decay, p= 0.002, t-test). Notably, all these parameters showed a sensitivity to CADASIL pathology close to 100%.

These encouraging results prompted us to investigate whether such fast fMRI data could serve to obtain potential discriminative biomarkers for the disease. We thus used the F1-score as a conservative measure of discrimination between the two samples, ranging from 50% (chance level) to 100% (perfect discrimination) and independent of t-test significance. For example, although the skewness and kurtosis differed significantly between the CADASIL and control groups, their F1-scores were found relatively low (67% and 68%, respectively). To look for higher F1-scores, we computed seven additional parameters to evaluate other features of responses. Figure 3d shows the F1-scores for all calculated parameters in the calcarine (the list of values for the other ROIs is presented in Table 1). We isolated the four parameters with the highest F1-scores (Fig.3e, besides the time to peak reported in 3c). The best-performing parameters achieved F1-scores of up to 82%. To assess the generalization power of the derived parameters, we estimated the corresponding effect sizes (Fig. 3d) with a mean Cohen’s *d* of 1.06 ± 0.17 (mean ± st.dev.). Remarkably, all seven parameters showed a sensitivity for CADASIL higher than 90%, and four of them even reached 100% (Table 1). We finally compared the response amplitudes of all ROIs. Although the amplitude in the BOLD signal varied more across experiments than subjects, no significant difference was detected between the CADASIL and control groups (Suppl Fig. 5 and Suppl. Table 2). Overall, our results emphasize that careful voxel selection within the different regions analyzed by fast fMRI is essential for detecting differences between CADASIL patients and controls, and may account for the inconsistent or null findings reported in previous studies of comparable cohorts.

**Table 1.** F1-score, sensitivity, and effect size of the 5 most informative fMRI parameters at 2 and 10-second stimulation in the Calcarine Sulcus, with the t-test selection, and in the visual cortex. Although the 5 parameters generally distinguished patients from control subjects, the selected ROI affected the statistical results.

| 2-sec stimulation |  | F1-score (%) |  |  | Sensitivity (%) |  |  | Effect Size |  |  |
| --- | --- | --- | --- | --- | --- | --- | --- | --- | --- | --- |
|  |  | Calcarine Sulcus | t-test selection | Vis. cortex | Calcarine Sulcus | t-test selection | Vis. cortex | Calcarine Sulcus | t-test selection | Vis. cortex |
| Area | AUC ratio | 80 | 79 | 76 | 100 | 83 | 89 | 1.21 | 0.88 | 0.91 |
| Duration | Zero-cross | 75 | 78 | 73 | 100 | 89 | 100 | 0.83 | 1.04 | 0.51 |
| Decay | Time half decay | 82 | 79 | 78 | 100 | 83 | 100 | 1.18 | 0.99 | 0.78 |
| Shape | Kurtosis Skewness Product | 67 | 67 | 67 | 94 | 94 | 94 | 0.03 | 0.30 | 0.19 |
|  | Full Width Half Max | 80 | 84 | 68 | 100 | 100 | 94 | 0.86 | 0.96 | 1.00 |
| Onset | Time half peak | 79 | 79 | 83 | 94 | 94 | 94 | 0.89 | 0.59 | 1.01 |
| Peak | Time to Peak | 81 | 78 | 78 | 94 | 89 | 89 | 1.13 | 0.58 | 0.59 |
| 10-sec stimulation |  | F1-score (%) |  |  | Sensitivity (%) |  |  | Effect Size |  |  |
|  |  | Calcarine sulcus | t-test selection | Vis. cortex | Calcarine Sulcus | t-test selection | Vis. cortex | Calcarine Sulcus | t-test selection | Vis. cortex |
| Area | AUC ratio | 80 | 79 | 82 | 82 | 88 | 82 | 1.11 | 1.02 | 1.25 |
| Duration | Zero-cross | 79 | 79 | 80 | 88 | 76 | 94 | 0.79 | 1.05 | 1.39 |
| Decay | Time half decay | 82 | 76 | 82 | 82 | 82 | 82 | 1.27 | 0.95 | 1.18 |
| Shape | Kurtosis Skewness Product | 80 | 78 | 85 | 94 | 82 | 100 | 1.40 | 1.17 | 1.27 |
|  | Full Width Half Max | 72 | 73 | 72 | 77 | 88 | 100 | 0.59 | 1.05 | 0.28 |
| Onset | Time half peak | 74 | 69 | 74 | 82 | 100 | 82 | 0.54 | 0.18 | 0.10 |
| Peak | Time to Peak | 79 | 69 | 71 | 100 | 100 | 100 | 1.30 | 0.23 | 0.14 |

Additional results further suggest that different features of hemodynamic alterations could be observed during both short and prolonged vascular responses in CADASIL. Indeed, we next examined also the responses to 10-second stimuli, which are expected to elicit stronger vascular responses and should be sufficient to saturate the response amplitude, based on our re-analysis of prior data (Suppl. Fig. 1). As with the 2-second stimuli, 10-second responses also showed a global slowdown of functional hyperemia in CADASIL patients when compared with controls (Fig. 4a). Note that one CADASIL patient could not be tested for 10-seconds stimulation because of technical issue of the MR scanner. Analyses within the calcarine area yielded results close to those obtained with the shorter stimuli (Fig. 4b, c). However, unlike the short-stimulus condition, the parameters derived from the t-test–based voxel selection showed no significant differences or only very weak effects (Table 1). By contrast, the selection based on the entire visual cortex produced more significant group differences with 10-second stimuli than with 2-second stimuli (Fig. 4b, c). This apparent inversion in the relative informational value of the t-test–selected voxels versus the anatomically defined visual cortex ROIs may reflect the engagement of additional components of neurovascular coupling during prolonged activation.

**Figure 4.**
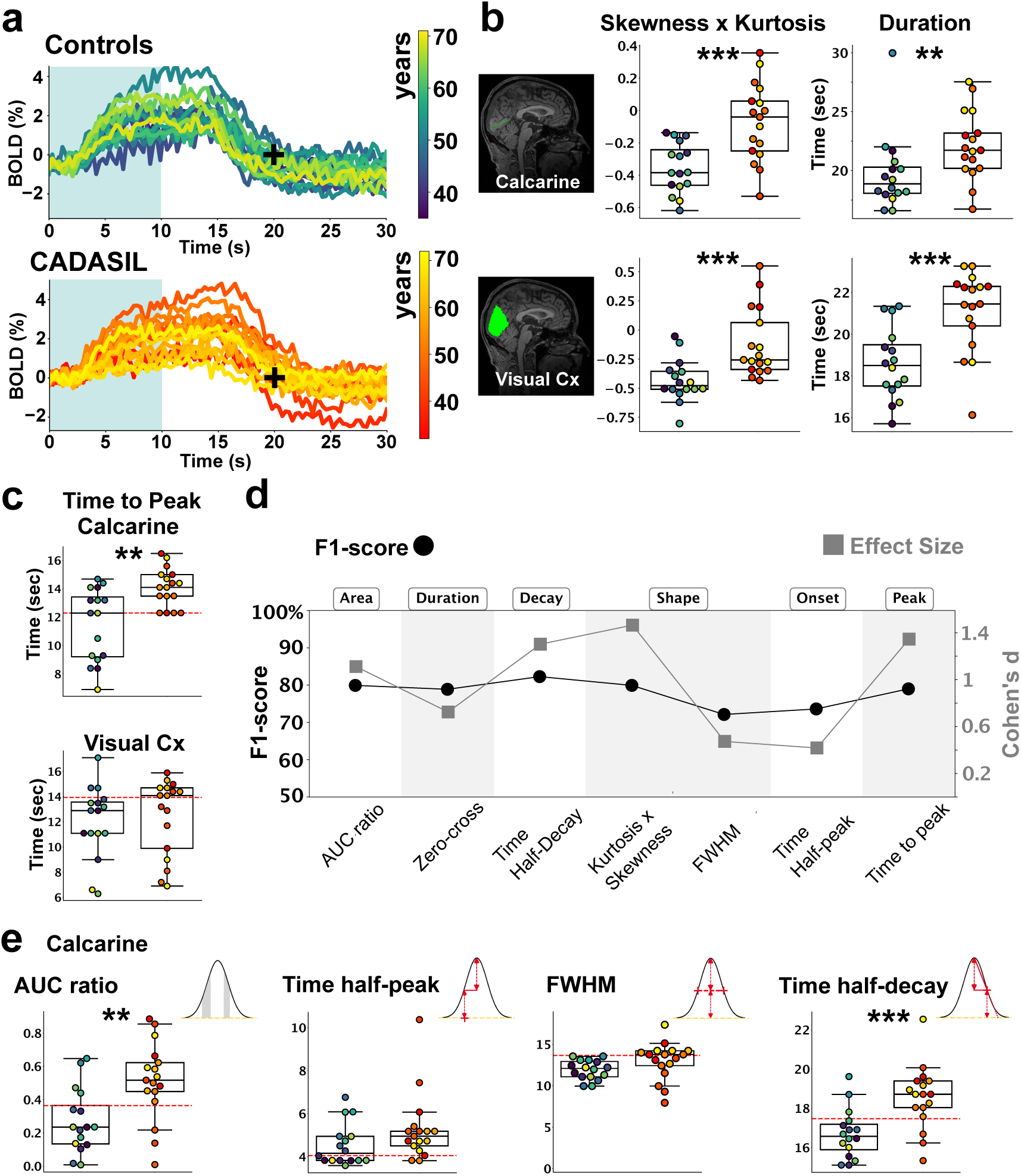
Long visual stimulation in CADASIL and Control subjects. (a) BOLD responses to 10 sec stimuli averaged in the calcarine for CADASIL and Control subjects, with color-coded subjects’ age. (b) The quantification of responses’ skewness x kurtosis and duration in calcarine and visual cortex shows a slower response in CADASIL than control subjects (Calcarine: skewness x kurtosis, p= 0.0004, duration, p= 0.006. Visual cortex: skewness x kurtosis, p= 0.0004, duration, p= 0.0006, t-test). (c) Time to peak of the BOLD response in calcarine, but not in visual cortex, further confirmed the slowdown for CADASIL’s responses versus Controls (Calcarine: p= 0.001. Visual cortex, p= 0.36, t-test). (d) F1-score and effect size values across 7 computed parameters accounting for different phases and features of the BOLD responses indicate a discrimination power in the order of 80%. Generalization assessment varied across the parameters, but it reached values close to 1.4 (Cohen’s d) in three parameters. (e) Selection of the best F1-score ranked parameters, the same as for the short responses (Calcarine: AUC ratio, p= 0.003, time half peak, p= 0.06, FWHM, p= 0.007, time half decay, p= 0.001, t-test).

For the calcarine ROI, we finally applied the previous multiparametric analysis to obtain a more straightforward comparison between short and long stimulus responses (Fig. 4d). We found high F1-scores and size effects (maximum values of 82% and 1.4, respectively), which were close to those calculated with short stimuli. The long stimulus parameters were also found to have high sensitivity, as illustrated by the time to peak reaching 100% sensitivity (Table 1). Finally, the analysis allowed us to isolate the same pool of parameters as those chosen for short stimuli (Fig. 4e).

Overall, our results demonstrate that CADASIL is associated with a consistent pattern of hemodynamic slowing across both short and prolonged visual stimulations, with differences in amplitude and spatial distribution depending on stimulus duration.

#### Further dissecting neurovascular coupling alterations in CADASIL

Because alterations were detected in both the rising and decay phases of the BOLD response in CADASIL, we asked whether these changes reflect a single dominant physiological disturbance or multiple independent processes. To indirectly address this question, we compared several response features (Fig. 5a) and quantified the correlations between the most discriminative related parameters across the two stimulus durations (Fig. 5b). The consistent and strong inter-feature correlations indicate that the slowing of the hemodynamic response is best explained by a common, or closely linked, underlying mechanism.

**Figure 5.**
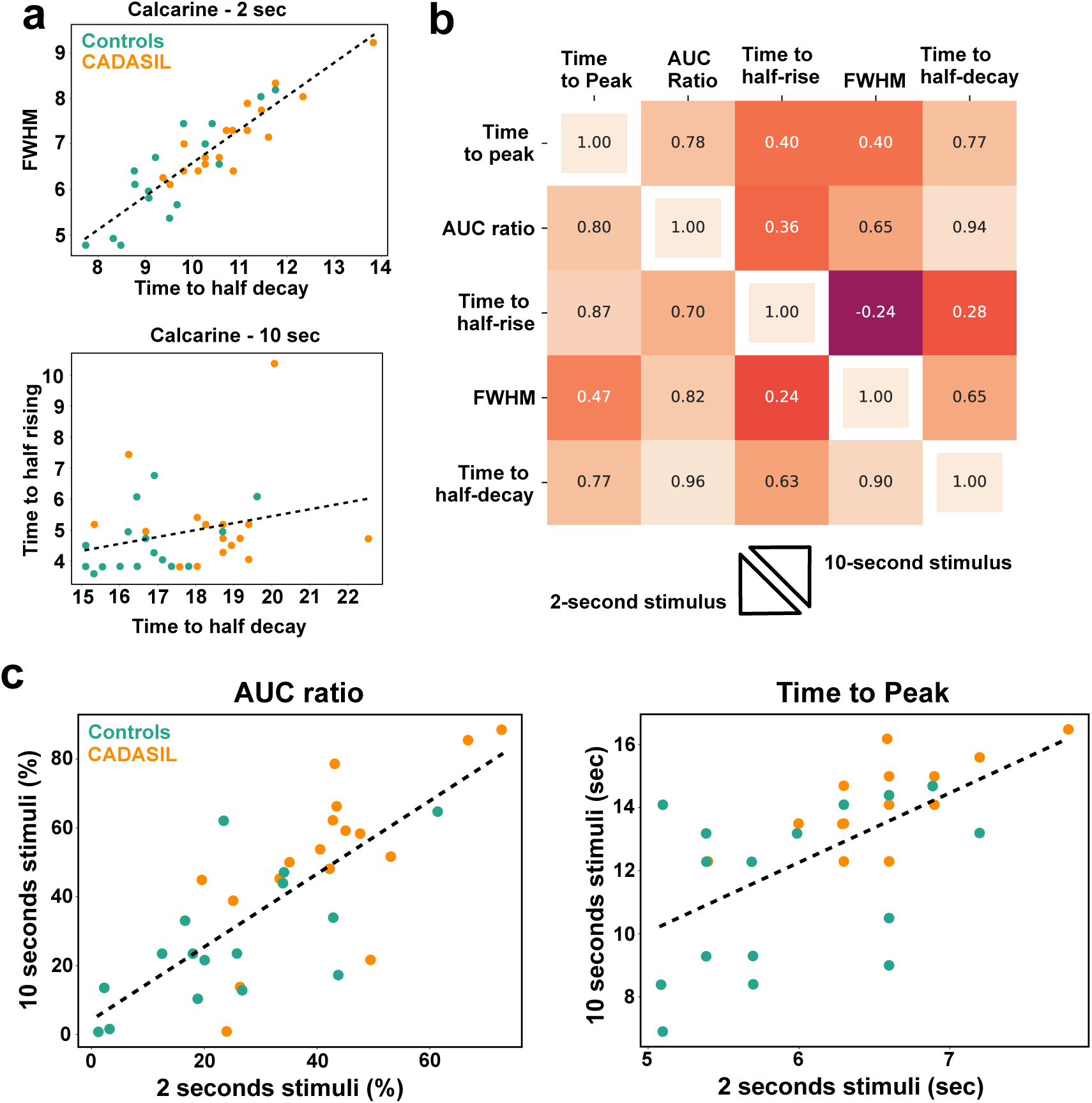
Considering multiple dynamic fMRI parameters does not improve F1-score. (a) Mixing parameters does not help to boost the CADASIL/Control prediction. In fact, most of the parameters are highly correlated for 2 and 10 sec responses. (b) Complete assessment of the correlation across all the computed parameters. Most of the tested pairs of parameters show a correlation over 70%. (c) mixing the values of the same parameter computed on 2 or 10 sec responses displayed a high correlation, hindering the possibility of boosting the prediction quality, as shown here for the AUC and the time to peak parameters.

We also compared the same parameter between 2 and 10-second stimuli (Fig. 5c) to assess whether short and prolonged responses might reflect distinct pathological processes. The strong correlations observed across stimulus durations provided no evidence for independent or multiple impaired components. These findings further indicate that combining dynamic parameters is unlikely to enhance discriminative performance, as they largely capture the same underlying abnormality.

As noted above, the BOLD amplitude must be interpreted with caution because of its intrinsic scan-to-scan variability (see also Suppl. Fig. 2). Nonetheless, the absence of any amplitude difference between CADASIL and control subjects for short stimuli across all tested ROIs (Suppl. Table 2) provides a stable reference (normalization factor) for interpreting the amplitudes measured during 10-second stimulations. Looking at the Calcarine or the t-test ROIs, we confirmed the lack of difference in the BOLD amplitude for the responses triggered by 10-second stimuli (Fig. 6a, b, and Suppl. Table 2). In contrast, we found a reduction of the amplitude of the BOLD response in CADASIL patients when the entire visual cortex was considered (Fig. 6c) although the number of significantly activated voxels upon the 10-sec stimulation did not really differ between CADASIL and control subjects (Mann-Whitney U-test, p = 0.31). Altogether, these findings suggest that, independently of the stimulus duration, the BOLD amplitude is preserved in the brain areas with the highest neuronal activation in CADASIL patients. Conversely, within the visual cortex, the hemodynamic response appears significantly smaller outside these areas, particularly, when the stimulation is prolonged.

**Figure 6.**
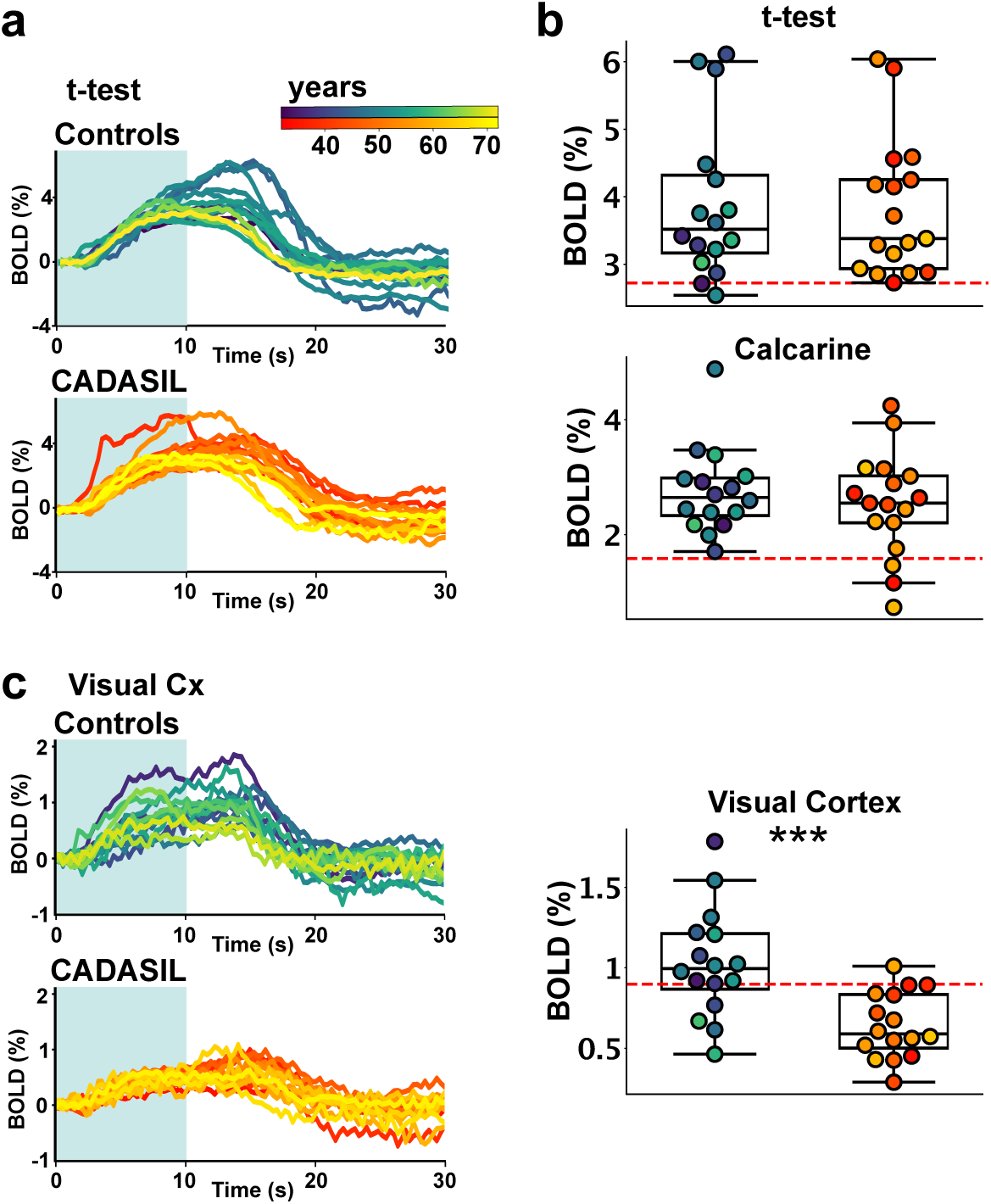
CADASIL patients have smaller BOLD responses in the visual cortex with 10 seconds stimulation. (a) Average BOLD responses to 10 sec visual stimulation from voxels selected as significantly activated with a t-test analysis (p= 0.93, t-test). (b) The amplitude of average responses from t-test-selected voxels or from the calcarine was not significantly different between CADASIL patients and control subjects (p= 0.43, t-test). (c) Average BOLD responses from voxels in the visual cortex was significantly lower in CADASIL than control subjects, as reported in the plot on the right (p= 0.0006, t-test).

Given the marked contrast in amplitude between strongly and weakly activated regions in CADASIL, we next examined whether this deficit was associated with slowing of the visually evoked response. When comparing the dynamic parameters measured in the calcarine with the amplitude extracted from the entire visual cortex, we found only weak correlations (Fig. 7a), suggesting that these alterations arise from partially independent pathological processes and supporting the view that neurovascular coupling is multifaceted in both health and disease. Owing to their limited correlation, the combination of whole–visual cortex BOLD amplitude and calcarine ROI time-to-peak produced an almost perfectly discriminative classifier distinguishing CADASIL patients from controls (Fig. 7a).

**Figure 7.**
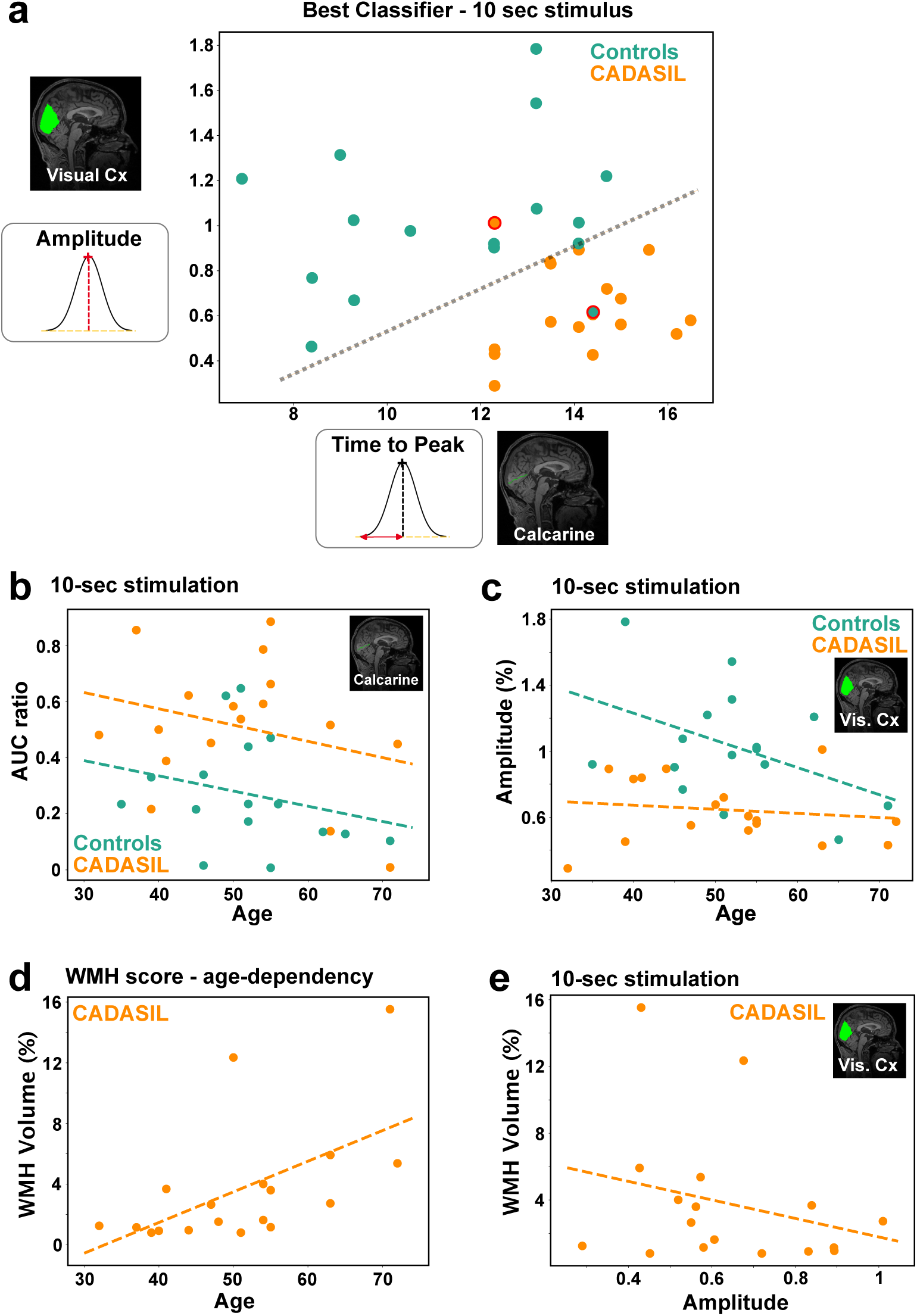
Relation of fMRI parameters, age and WMHs. (a) The best classifier for CADASIL patients and control subjects was obtained by combining parameters computed from 10-sec BOLD responses. The amplitude from the entire visual cortex and the Time to peak from the calcarine almost perfectly separated the two groups of subjects. One false positive and one false negative subject were misclassified. (b) The vast majority of fMRI-derived parameters showed no age dependence, regardless of stimulus duration, as exemplified by the AUC ratio extracted from the calcarine during 10-second stimulation (Control subjects: p= 0.34, CADASIL patients, p= 0.36, F-test). (c) The BOLD amplitude evoked by 10-second stimuli across the entire visual cortex displayed a decreasing trend with age in control subjects (Control subjects: p= 0.12, F-test), while it was flat in CADASIL patients, indicating a clear early aging deficit in the patients (CADASIL patients: p= 0.61, F-test). (d) The automatic quantification of the volumes of the White Matter Hyperintensities (WMHs), normalized by the brain volume, in CADASIL patients showed an increasing trend with age (p= 0.016, F-test). (e) No clear correlation between WMHs and one of the most discriminative fMRI parameters, the amplitude of the response extracted from the visual cortex (p= 0.31, F-test).

Leveraging the wide lifespan of our study participants, we also investigated the age dependency of the response dynamics and amplitude in both healthy subjects and CADASIL patients. Among all parameters with the highest F1-score extracted from BOLD fMRI data obtained in the Calcarine Sulcus, only the time-to-peak was found to be related to age in healthy individuals but not in patients (Suppl. Table 3 and Fig. 7b). Interestingly, a trend toward a decrease of BOLD amplitude along aging was also observed in healthy individuals for the 10-second stimulation in the Visual Cortex (Fig. 7c). No age effect was detected on the response amplitude in patients who presented since early stage with values close to those measured in the oldest individuals of the control group. The volume of white Matter Hyperintensities (WMHs) normalized to the intracranial cavity in each individual was correlated to age in CADASIL patients (Fig. 7d). However, no correlation was detected between WMH volumes and the different computed fMRI parameters obtained from the Calcarine (Suppl. Table 4) or 10-second stimuli amplitude extracted from the visual cortex (Fig. 7e).

## Discussion

In the present study, we explored the potential of fast BOLD fMRI to assess neurovascular coupling alterations using a 3T MR scanner commonly used in clinical practice. We tested a short visual stimulation sequence that allows functional hyperemia responses to be explored with sufficient time for complete recovery. The effects of two stimulation durations for measuring a short and a more demanding vascular response, which are presumably mediated by distinct cell types and molecular mechanisms^24–26^, were specifically investigated. The results were obtained at the individual level to measure the discriminatory power of multiple extracted parameters, in accordance with previous recommendations.^22^

We first demonstrated that our approach enables single-repetition analyses of short visual stimuli at standard fMRI spatial resolution. Applying fast BOLD fMRI in a paradigmatic monogenic cSVD, CADASIL, we detected a disease-related slowing of visually evoked responses that varied markedly with the level of topographic focus used for voxel analysis. Previous fMRI studies in CADASIL reported either increased BOLD amplitudes^13^ or the opposite.^21,22,27^ Here, we found preserved amplitudes when averaging responses within the calcarine sulcus, but a marked decrease when responses were averaged across the entire visual cortex—an effect that was particularly pronounced for longer stimulations—thereby partially reconciling previous discrepant findings.

Our study illustrates the critical importance of ROI definition, when extracting stimulus-evoked fMRI time courses, particularly when aiming to characterize the detailed shape of the hemodynamic response. This step is most often overlooked, with ROIs frequently chosen arbitrarily and very rare studies systematically assessing how different topographic definitions could dramatically influence the results^28^. We selected ROIs to avoid circular analysis issues^28,29^ using either anatomy-driven definitions (e.g., calcarine sulcus, visual cortex) or independent statistical contrasts (t-tests) that do not impose explicit assumptions on HRF shape or dynamics.

In addition, we used an AI-based approach to localize and delineate the calcarine sulcus in each subject and manually separated the calcarine from the peri-calcarine region to refine ROI definition. This procedure revealed that ROI selection markedly influences the response amplitude and signal-to-noise ratio, while leaving the temporal dynamics of the BOLD response largely unchanged.

When examining these ROIs in relation to the pathology, we observed marked differences in discrimination performance across both temporal (dynamics) and spatial dimensions. Notably, the calcarine cortex provided the most robust and reliable assessment of BOLD response slowing in CADASIL patients across both 2- and 10-second visual stimulation paradigms. In contrast, analyses at the whole-visual-cortex level revealed a significantly reduced BOLD amplitude in CADASIL, but only in response to longer stimuli. Together, these spatial and temporal findings support the working hypothesis that CADASIL can be probed through two distinct neurovascular coupling regimes, depending on stimulus duration.

Our results are consistent with preclinical investigations in CADASIL mouse models showing a defect in vascular backpropagation signaling during neurovascular coupling^30–35^. Although the reduced spatial spread of large activations in CADASIL patients could, in theory, arise from diminished neural and vascular signaling, the markedly slow vascular dynamics observed in CADASIL are more consistent with intrinsic vessel-wall dysfunction, in line with alterations of smooth muscle cells^28^ and pericytes in such conditions^29^. This interpretation is also consistent with the similar slowdown of vascular dynamics detected in the skin and retinal microvascular reactivity^36,37^.

Fast BOLD fMRI investigations helped us dissect subtle dynamic differences in vascular responses, despite BOLD contrast at 3T is expected to arise mostly from signal modifications occurring at the venous level^6^. This unexpected dynamic sensitivity of fast BOLD fMRI in probing neurovascular coupling may open avenues for research or diagnostic applications inaccessible at standard fMRI temporal resolution. Furthermore, such a fast fMRI approach appears feasible using commercial MRI scanners and offers a high signal-to-noise ratio, potentially allowing its use with 1.5 T scanners. Previous fMRI studies in CADASIL patients did not show significant modifications in BOLD response dynamics^13,27^. This suggests that fast fMRI with high temporal resolution was needed to capture the response slowing observed here in CADASIL. Alternatively, previous paradigms may not have allowed the BOLD signal to return to baseline between successive stimuli, potentially inducing a ceiling effect in neurovascular coupling and masking dynamic differences.

In healthy individuals, the amplitude of the BOLD response in the visual cortex declined with age, while its temporal dynamics remained largely preserved—a pattern consistent with the well-documented age-related reduction in cerebrovascular reactivity. In contrast, CADASIL patients exhibited no age dependency in BOLD amplitude, suggesting that their cerebral vessels resemble those of older adults even at early disease stages, prior to the onset of cognitive decline or disability. This aligns with recent findings from retinal adaptive optics imaging^37^. Given the pronounced differences in fMRI responses between patients and controls, we extended our analysis beyond traditional group-level statistics to evaluate biomarker performance using metrics such as sensitivity and the F1-score. Notably, F1-scores reached 80% for many fMRI-derived parameters, independent of stimulus duration, with some parameters from the 2-second stimulation data achieving 100% sensitivity. Effect sizes for the computed parameters frequently exceeded 1 (e.g., 1.4 for 10-second stimulation), indicating robust group differences within our cohort. While the limited sample size precludes broad generalizability, the careful selection of patients with isolated cSVD pathology—excluding common comorbidities—enhances the internal validity of these findings and provides a foundation for future studies in larger, more diverse cSVD populations.

Finally, our data highlight the importance of probing neurovascular coupling using both short and long stimulations, which engage distinct neurovascular regimes—a brief activation for 2-second stimuli and a stronger, more sustained activation for 10-second stimuli. The slowdown of short and prolonged BOLD responses in CADASIL patients implicates defective blood-vessel dynamics even under mild activation. In such SVD, this deficit does not recover when the vasculature is solicited by a more demanding activation task. Moreover, the marked amplitude differences observed with 10-second stimuli are unlikely to be explained by inter-examination variability in the BOLD signal-to-noise, given the absence of any amplitude difference for 2-second stimuli within the same ROI.

We propose that this dual-stimulus framework should become a standard methodological approach for BOLD-fMRI studies in cSVD.

## Acknowledgments

This work was supported by the RHU Treat-SVD grant (ANR-16-RHUS-0004) and the ANR PRC 2023 grant fMRI-BioSVD, awarded to H.C., and to H.C. and D.B., respectively.

It was also conducted within the CERVCO reference center, in collaboration with the Institut Hospitalo-Universitaire Vascular Brain Institute (IHU-VHBI), supported by the French National Research Agency (ANR-23-IAHU-0001) as part of the “France 2030” program. We are also grateful to the Association de Recherche Neurovasculaire (ARNEVA) at Lariboisière and to the CADASIL France patient association for their continued support.

We extend our sincere gratitude to Mr. Mohamed Saichi for the segmentation of white matter lesions in the patients, and to Dr. Christian Giroux for his invaluable assistance in patient recruitment and enrollment.

During the preparation of this work, the authors used Grammarly in order to check proofreading. After using this tool/service, the authors reviewed and edited the content as needed and take(s) full responsibility for the content of the published article.

This study has been registered as a clinical trial: https://clinicaltrials.gov/study/NCT04036084?cond=cadasil&viewType=Card&page=2&rank=15

---

**Suppl. 1:**
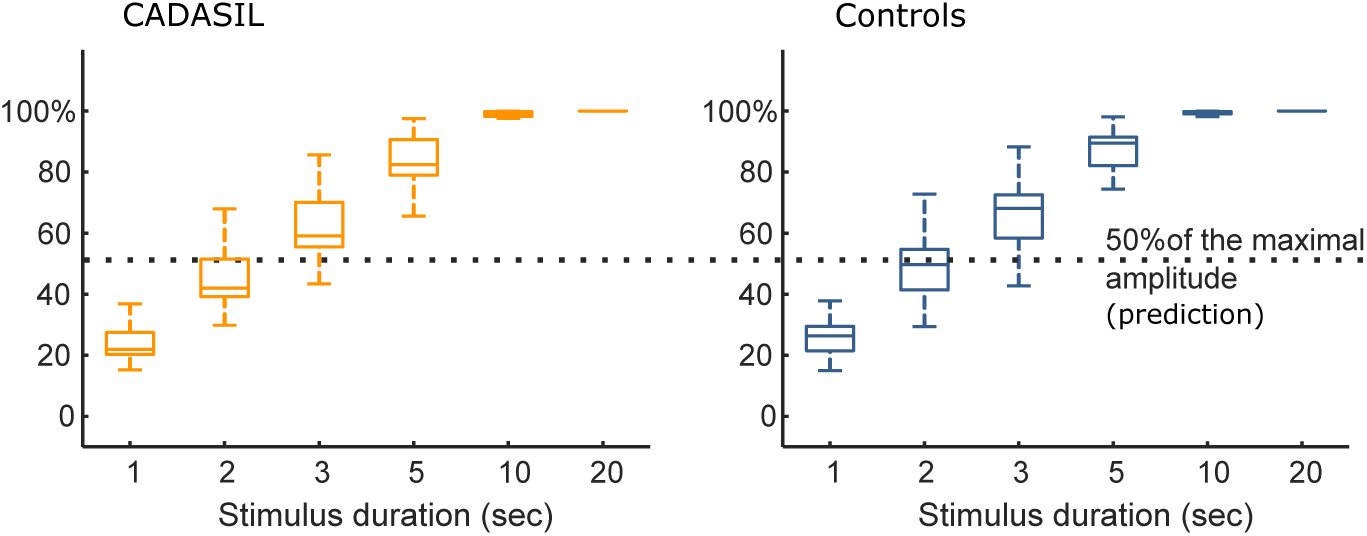
Convolution-based prediction of the visually stimulated BOLD response shortening the stimulus duration indicated that, for both CADASIL patients and control subjects, we expect to get the maximal amplitude and one-half of it with 10-second and 2-second stimuli, respectively. This led us to select these durations to study two neurovascular regimes.

**Suppl. 2:**
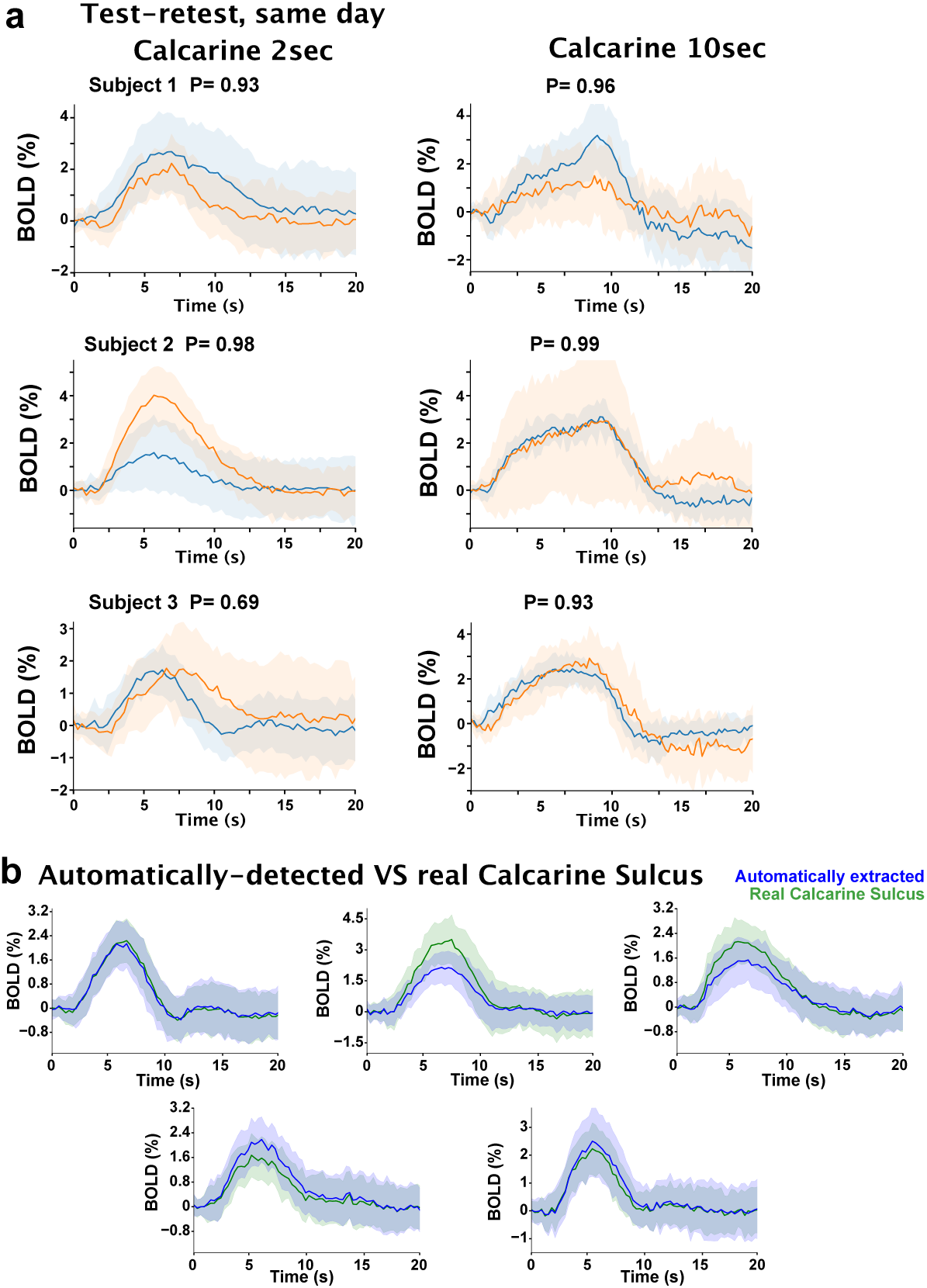
(a) Examples of test-retest BOLD responses within the same day on healthy subjects, for 2 and 10 seconds stimulations. (b) Examples of BOLD responses using the Calcarine + Pericalcarine sulci automatically segmented with BrainVISA Morphologist and the responses from the Calcarine Sulcus manually isolated from the Pericalcarine.

**Suppl. 3:**
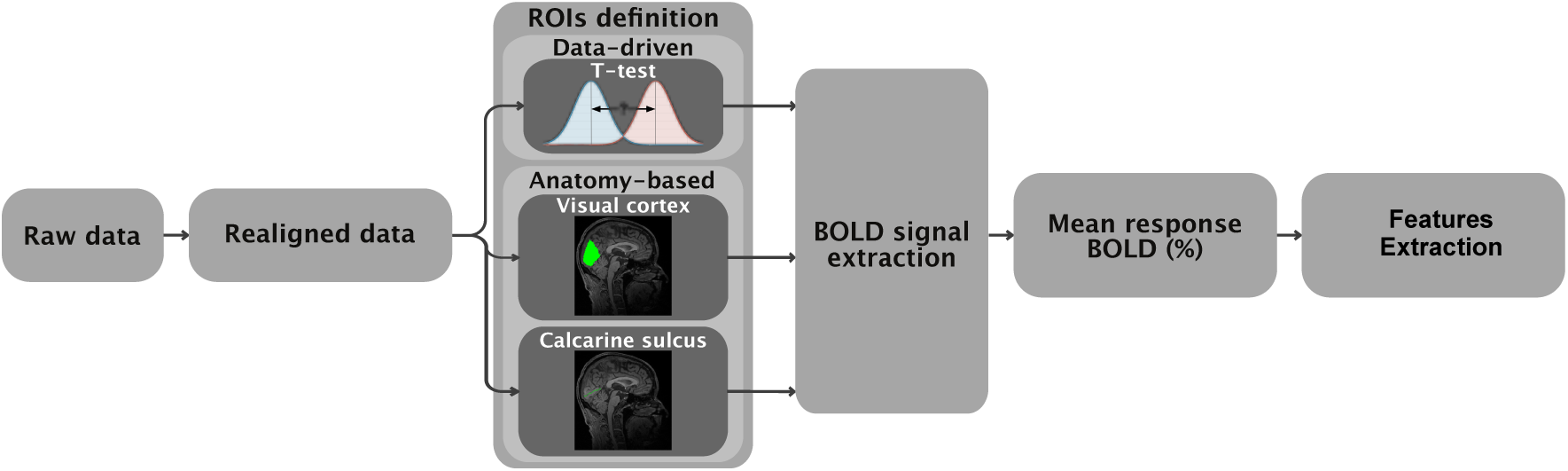
Schematics of the analytical pipeline used to extract features for the entire study. Thanks to the high temporal resolution of our data, we avoided the slice-timing correction.

**Suppl. 4:**
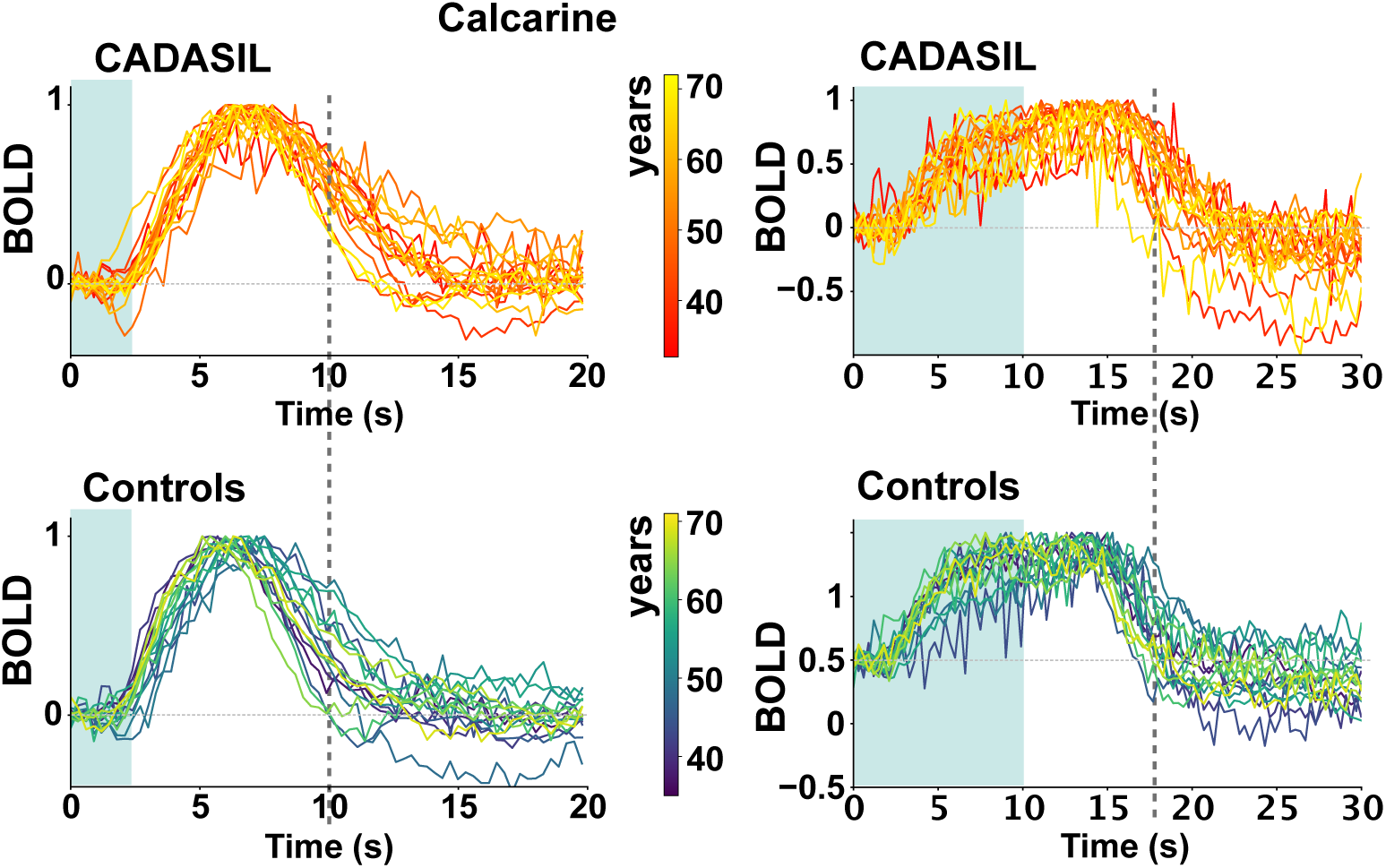
Normalized 2 and 10-seconds visually evoked BOLD responses for each subject, as reported in Fig. 3a and 4a, respectively. The difference in CADASIL dynamics is evident.

**Suppl. 5:**
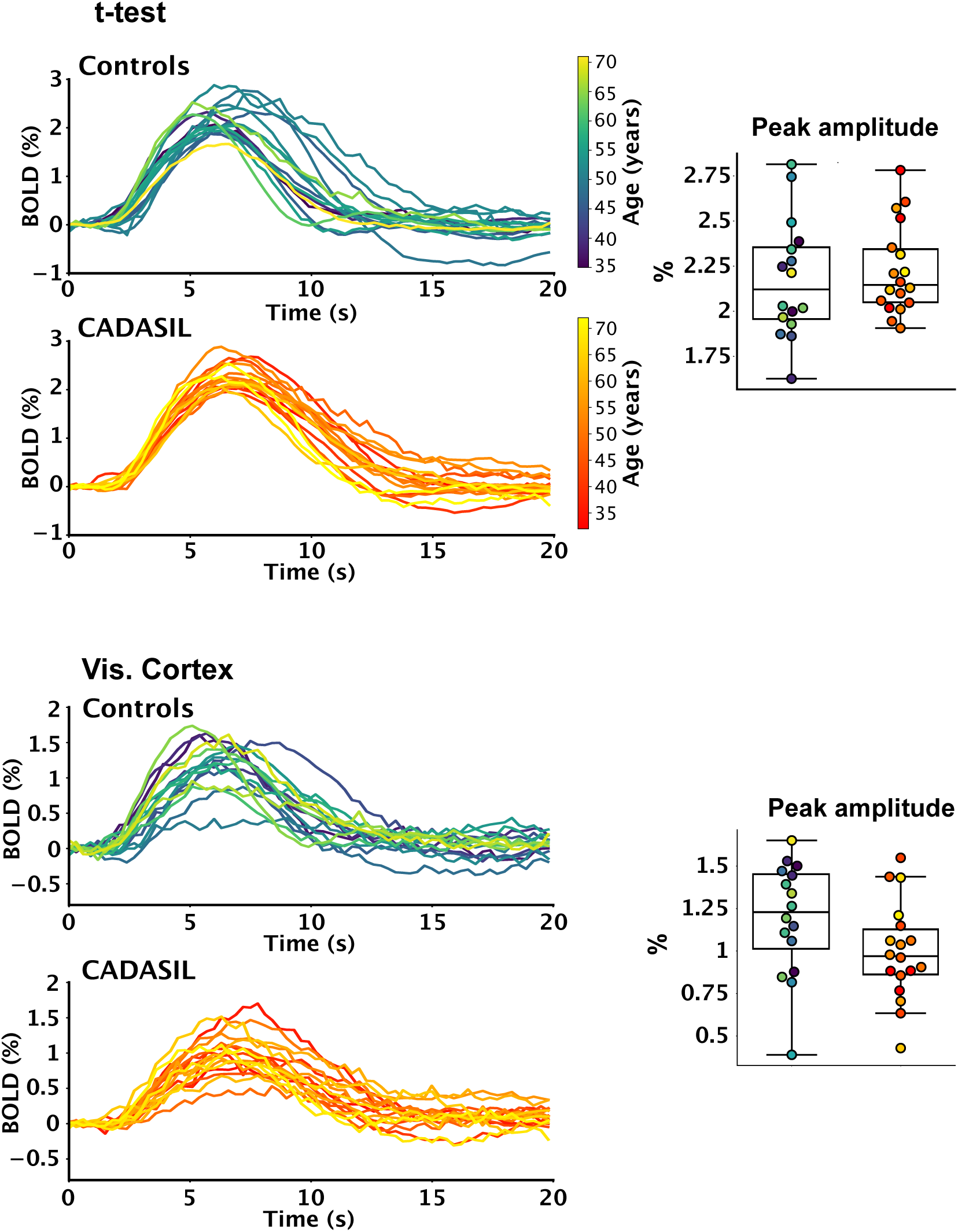
BOLD responses upon 2-second stimulation averaged on t-test-selected voxels or within the entire visual cortex. No difference was found in the amplitude between CADASIL and control subjects.

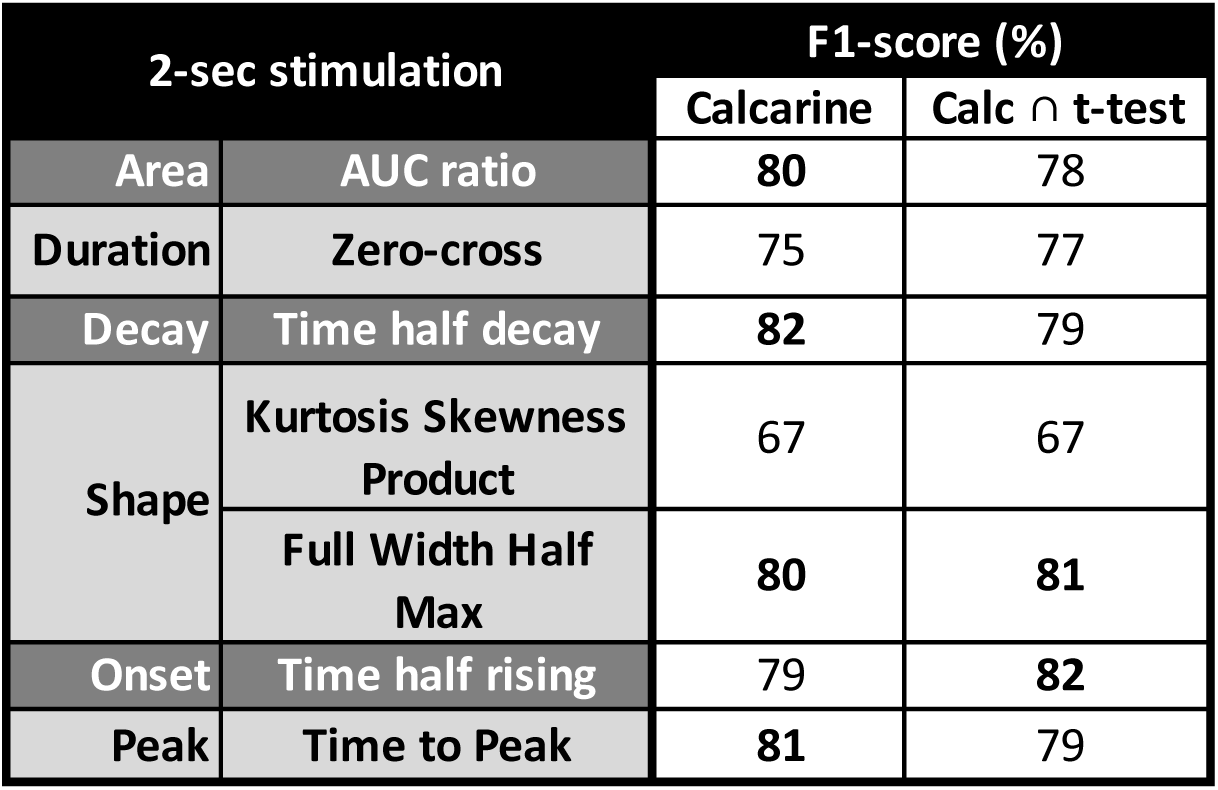

| Amplitude | p-value |  |
| --- | --- | --- |
|  | 2-sec stim. | 10-sec stim. |
| Calcarine | 0.35 | 0.43 |
| Visual Cortex | 0.08 | **0.00056** |
| t-test | 0.63 | 0.93 |

| Age correlation<br>p-values in the<br>Calcarine Sulcus | 2-sec stimulation |  | 10-sec stimulation |  |
| --- | --- | --- | --- | --- |
|  | Control<br>subjects | CADASIL<br>patients | Control<br>subjects | CADASIL<br>patients |
| AUC ratio | 0.94 | 0.18 | 0.34 | 0.26 |
| Time half decay | 0.94 | 0.63 | 0.20 | 0.25 |
| Full Width Half Max | 0.92 | 0.17 | 0.41 | 0.24 |
| Time half rising | 0.95 | 0.63 | 0.42 | 0.60 |
| Time to Peak | 0.70 | 0.63 | **0.02** | 1.00 |

| **CADASIL WMH-volume (%)**<br>**correlation with fMRI parameters in**<br>**Calcarine Sulcus (p-values)** | **2-seconds**<br>**stimuli** | **10-seconds**<br>**stimuli** |
| --- | --- | --- |
| **AUC ratio** | 0.17 | 0.11 |
| **Time half decay** | 0.14 | 0.21 |
| **Full Width Half Max** | 0.09 | 0.65 |
| **Time half rising** | 0.73 | 0.53 |
| **Time to Peak** | 0.69 | 0.59 |

